# mGlu_2_ and mGlu_3_ receptor negative allosteric modulators attenuate the interoceptive effects of alcohol in male and female rats

**DOI:** 10.1101/2022.04.16.488559

**Authors:** Ryan E. Tyler, Kalynn Van Voorhies, Bruce E. Blough, Antonio Landavazo, Joyce Besheer

## Abstract

**Rationale:** The subjective effects of alcohol are associated with alcohol use disorder (AUD) vulnerability and treatment outcomes. The interoceptive effects of alcohol are part of these subjective effects and can be measured in animal models using drug discrimination procedures. The newly developed mGlu_2_ and mGlu_3_ negative allosteric modulators (NAMs) are potential therapeutics for AUD and may alter interoceptive sensitivity to alcohol.

**Objectives:** To determine the effects of mGlu_2_ and mGlu_3_ NAMs on the interoceptive effects of alcohol in rats.

**Methods:** Long-Evans rats were trained to discriminate the interoceptive stimulus effects of alcohol (2.0 g/kg, i.g.) from water using both operant (males only) and Pavlovian (male and female) drug discrimination techniques. Following acquisition training, an alcohol dose-response (0, 0.5, 1.0, 2.0 g/kg) experiment was conducted to confirm stimulus control over behavior. Next, to test the involvement of mGlu_2_ and mGlu_3_, rats were pretreated with the mGlu_2_-NAM (VU6001966; 0, 3, 6, 12 mg/kg, i.p.) or the mGlu_3_-NAM (VU6010572; 0, 3, 6, 12 mg/kg, i.p.) before alcohol administration (2.0 g/kg, i.g.).

**Results:** In Pavlovian discrimination, male rats showed greater interoceptive sensitivity to 1.0 and 2.0 g/kg alcohol compared to female rats. Both mGlu_2_-NAM and mGlu_3_-NAM attenuated the interoceptive effects of alcohol in male and female rats using Pavlovian and operant discrimination. There may be a potential sex difference in response to the mGlu_2_-NAM at the highest dose tested.

**Conclusions:** Male rats may be more sensitive to the interoceptive effects of 2.0 g/kg alcohol training dose compared to female rats. Both mGlu_2_ and mGlu_3_ NAM attenuate the interoceptive effects of alcohol in male and female rats. Sex differences in mGlu_2_-NAM sensitivity were observed. These drugs may have potential for treatment of AUD in part by blunting the subjective effects of alcohol.

## Introduction

Alcohol use disorder is a chronic, relapsing disease associated with high morbidity and mortality [1]. The subjective effects of alcohol are an important factor in susceptibility and other outcomes for alcohol use disorder [2]. A recent meta-analysis found that laboratory measures of the subjective effects of alcohol (stimulation, sedation, and craving) are associated with clinical outcomes for AUD pharmacotherapies – i.e., medications that reduced stimulation, sedation, and craving in a laboratory study were associated with better clinical outcomes [3]. Furthermore, studies show that people with greater sensitivity to the stimulatory effects of alcohol and lower sensitivity to the sedative effects of alcohol are more likely to engage in binge drinking and to develop an alcohol use disorder [4–6]. Part of the subjective effects of alcohol include the interoceptive effects, which refer to the internal sense or perception of the body [7]. All drugs that have the potential to produce substance use disorder or addiction, including alcohol, produce distinct interoceptive effects that can influence subsequent drug taking by signaling satiety or driving more intake [7, 8]. As such, the mechanisms underlying the interoceptive processing of alcohol are an important and clinically relevant research avenue. Furthermore, drugs that modulate the interoceptive/subjective effects of alcohol may have potential as a therapeutic for AUD.

Metabotropic glutamate (mGlu) receptors are G protein-coupled receptors (GPCRs) activated by the neurotransmitter glutamate. There are three groups of mGlu receptors, Group I (mGlu_1_ and mGlu_5_), Group II (mGlu_2_ and mGlu_3_), and Group III (mGlu_4_, mGlu_6_, mGlu_7_, mGlu_8_). Group I receptors are predominantly expressed at the postsynaptic site and are coupled to Gq proteins, whereas Group II and III receptors are considered mostly presynaptic and coupled to Gi proteins [9, 10]. mGlu receptors are promising targets for their potential in the treatment of alcohol use disorder (AUD), with both preclinical and clinical studies implicating these receptors in alcohol-related adaptations and AUD [9, 11–15]. Numerous studies have shown that manipulation of mGlu receptors are effective at reducing drinking and reinstatement of alcohol seeking [16–19], as well as being involved in the interoceptive effects of alcohol [20–23]. In humans, AUD is associated with an increase in mGlu_5_ radioligand binding using positron emission tomography [24]. Furthermore, single nucleotide polymorphisms (SNPs) of the mGlu_3_ gene (*Grm3*) have been identified as highly associated with AUD in a Genome Wide Association Study (GWAS) [25]. Prior studies from our lab show that mGlu_2/3_ receptor agonism and antagonism disrupt the interoceptive effects of alcohol when using a 1.0 g/kg alcohol training dose [21, 22]. However, at the time, pharmacological limitations did not allow for the evaluation of the specific role of mGlu_2_ vs. mGlu_3_ receptors. This obstacle has recently been overcome by the development of highly selective mGlu_2_ (VU6001966) and mGlu_3_ (VU6010572) negative allosteric modulators (NAMs) [26, 27]. Additionally, allosteric modulators of mGlu receptors show particular promise for novel targets of AUD. Both NAMs show blood-brain barrier permeability, high specificity for their respective receptors and both in vitro and in vivo efficacy [26–28]. In vivo effects include acute anti-depressant-like effects [26, 27] and prophylactic effects on measures of anxiety and arousal-like behaviors for the mGlu_3_-NAM [28]. The present work are the first studies to evaluate these drugs on the interoceptive stimulus effects of alcohol in rats.

Using drug discrimination techniques, animals can be trained to discriminate the interoceptive effects of alcohol from vehicle (e.g., saline, water) using operant and Pavlovian drug discrimination techniques [7, 20–23, 29–38]. In the present study the functional role of mGlu_2_ and mGlu_3_ receptors in the interoceptive effects of alcohol (2.0 g/kg) were assessed using both operant and Pavlovian discrimination procedures. The alcohol training dose of 2.0 g/kg was chosen because the contribution of glutamatergic signaling in the discriminative stimulus of alcohol appears to be greater at doses higher than 1.5 g/kg [39, 40]. As previous work has shown that mGlu_2/3_ antagonism attenuated the discriminative stimulus effects of alcohol in previous work [22], we hypothesized that negative allosteric modulation of mGlu_2_ and mGlu_3_ using VU6001966 and VU6010572, respectively would also attenuate the interoceptive effects of alcohol. Mechanistically we hypothesized that this consequence would occur through increasing excitatory drive [41], and therefore blunting the ability of alcohol to decrease excitation in the brain [7]. While the operant discrimination experiment assessed the NAMs in males only, the Pavlovian discrimination experiment assessed the effects in males and females and used a separate cohort of animals for each dose-response drug test to obviate potential interference with prior drug treatment.

## Methods

### Animals

Long Evans rats (Envigo, Indianapolis, IN) were used for all experiments. Rats arrived at 7 weeks and were housed individually. The vivarium was maintained on a 12-hr light/dark cycle and all experiments were conducted during the light cycle. All rats were handled for at least 1LJminute daily for 1 week prior to beginning discrimination training. Rats were fed daily to maintain body weight at 85% of free-feeding weight. Rats had ad libitum access to water. Rats were under continuous care and monitoring by veterinary staff from the Division of Comparative Medicine at UNCLJChapel Hill. All procedures were conducted in accordance with the NIH Guide to Care and Use of Laboratory Animals and institutional guidelines.

### Drugs

Alcohol (95% (v/v), Pharmaco-AAPER, Shelbyville, KY) was diluted with tap water to make a 20% alcohol solution for all experiments. mGlu_2_-NAM (VU6001966) was purchased from Tocris Bioscience (Bristol, UK). mGlu_3_-NAM (VU6010572) was synthesized by the Research Triangle Institute (RTI). Both drugs were dissolved completely in a 45% β-cyclodextrin vehicle at a concentration of 3, 6 or 12 mg/mL.

### Experiment 1: Operant Drug Discrimination: Effect of mGlu_2_ and mGlu_3_ NAM on the interoceptive effects of alcohol

#### Operant Drug Discrimination: Apparatus

The chambers (Med Associates, Georgia, VT; measuring 31 × 32 × 24 cm) used for drug discrimination training and testing have been detailed elsewhere [31, 35]. Chambers were inside of a sound-attenuating cubicle with an exhaust fan. Briefly, two retractable levers were on each side (left or right) of a liquid dipper receptacle on the right-side wall of the chamber. One cue light was located above each lever. Completion of a fixed ratio 10 (FR10) schedule of reinforcement on the appropriate lever activated a dipper that presented 0.1 mL of sucrose (10% w/v) for 4 seconds. Med Associates program was used to control sessions and record data.

#### Operant Drug Discrimination: Training

First rats underwent lever press training as detailed in [32]. Discrimination training began once responding on the FR10 schedule was stable (<10% daily variation in the total number of responses). Training sessions were similar to our previous work [20, 22, 23, 29, 31, 32, 34–36, 42]. Briefly, training sessions took place each day from Monday-Friday. Rats (N=9 males) were administered alcohol (2.0 g/kg, i.g.) or water (i.g.) and immediately placed in the chamber for a 20-min timeout period before the session started. Once the session started, both the right and left levers were introduced into the chambers and the house light was illuminated to indicate the beginning of the training session. During water training sessions, completion of FR10 on the water-appropriate lever (e.g., right lever) resulted in sucrose presentation and the illumination of the cue light above the lever. In contrast, on alcohol training sessions, completion of FR10 on the alcohol-appropriate lever (e.g., left lever) resulted in sucrose presentation and the illumination of the cue light above the lever. For each session, responding on the inappropriate lever was obtained but did not produce a programmed consequence. Alcohol- and water-appropriate levers were counterbalanced across rats. Training days varied on a double alternating schedule (alcohol, alcohol, water, water,…). Testing began when the following performance criteria were met: ≥ 80% appropriate lever responses for at least 8 out of 10 consecutive days for the first reinforcer and for the entire session.

#### Operant Drug Discrimination: Testing

Test sessions were identical to training sessions except that they were 2-min in duration and FR10 on either lever resulted in sucrose delivery to prevent lever selection bias and to allow for quantification of response rate. Identical to training sessions, a 20-min delay period elapsed from alcohol administration to the start of the test session (2 min).

Prior to testing, an alcohol dose response curve (0.0, 0.5, 1.0, 2.0 g/kg, i.g.) was generated to confirm stimulus control of behavior (N=9). N=6 rats were used for initial testing of the mGlu_3_-NAM. Next, the mGlu_2_-NAM was tested in the same rats but 2 were replaced, due to performance attrition during intervening training sessions, again for an N=6. For the NAM drug testing, rats were injected with the mGlu_3_-NAM VU6010572 (0, 3, 6 mg/kg, i.p., N=6) or mGlu_2_-NAM VU6001966 (0, 3, 6 mg/kg, i.p., N=6) 25 mins before alcohol (2.0 g/kg, i.g.) administration, such that 45 mins elapsed from the time the drug was administered to the start of the operant test session (25-min drug pretreatment + 20-min alcohol pretreatment). For each drug, doses administered were counterbalanced across days and all rats received all doses. Drug testing days were interspersed with training sessions. Because 6 mg/kg VU6010572 (mGlu_3_-NAM) did not produce an effect on alcohol-appropriate lever responding or response rate, a follow-up test was conducted using a higher dose (12 mg/kg) of mGlu_3_-NAM.

### Experiment 2: Pavlovian Drug Discrimination: Effect of mGlu_2_ and mGlu_3_ NAM on the interoceptive effects of alcohol

This experiment was conducted in both males and females using a different group of rats for each drug tested (mGlu_2_ and mGlu_3_ NAM). Because the 12 mg/kg dose was tested post-hoc in Experiment 1, the 12 mg/kg dose was included in the dose-response testing. Starting sample sizes for each group were as follows: male mGlu_2_-NAM (N=9); female mGlu_2_-NAM (N=8), male mGlu_3_-NAM (N=10), and female mGlu_3_-NAM (N=15).

#### Pavlovian Drug Discrimination: Apparatus

This apparatus is described in [30] and the chambers were identical to those used for operant discrimination (see above) with the exception that the levers were always retracted. On one side of the chamber, cue lights were located on either side of a liquid dipper receptacle and a photobeam detector was used to measure head entries into the liquid receptacle. When activated, the dipper was raised for 4 seconds and presented 0.1 mL of sucrose (26% w/v). Chambers were also outfitted with infrared photobeams (which divided the chamber into four parallel zones) to measure locomotor activity during sessions (number of beam breaks).

#### Pavlovian Drug Discrimination: Training

Training began with sucrose access training that is identical to the lab’s previous work [30, 34, 37, 43, 44]. Training started with three 50-min sessions in which sucrose (26 %) was presented randomly throughout the session to train the rats to approach the liquid receptacle. The probability of sucrose presentation decreased from the first to the last session and by the last 10 min of the final session rats received approximately 0.75 sucrose presentations/min. Following sucrose access training, the discrimination training began. Similar to the operant training procedure, training sessions were conducted 5 days per week (M-F) during which alcohol (2.0 g/kg) or water was delivered i.g. before the start of the training session. A 20-min timeout period from alcohol/water administration to the start of the training session (identical to the operant discrimination training in Experiment 1) and during this time, no cue lights were illuminated, no sucrose was presented, and head entries into the liquid receptacle were not recorded. After this delay, the 15-min session started. During alcohol training sessions, the offset of each of the 15-second cue light presentations (10 presentations in total) was followed by presentation of sucrose (26%, 0.1 mL, 4 sec). In contrast, during water training sessions, sucrose was not delivered following the offset of the cue light presentations. Like operant discrimination (Experiment 1), training sessions (days) occurred on a double alternating schedule (A, A, W, W…). Using the mean discrimination score from the previous 2 water and 2 alcohol sessions, the [alcohol session score – water session score] had to be ≥ 3 to meet criteria for testing [45–47]. Testing started once the criteria were met.

#### Pavlovian Drug Discrimination: Testing

Test sessions consisted of the standard 20-min timeout period, and then a single 15-s cue light presentation. No sucrose was delivered following the offset of the light presentation. Prior to testing an alcohol dose response curve (0.0, 0.5, 1.0, 2.0 g/kg, i.g.) was generated to confirm stimulus control. Identical to operant drug discrimination testing, the mGlu_2_ or mGlu_3_ NAM was administered 45-min before the start of the test session and 25 min before the alcohol administration (2.0 g/kg, i.g.). All 4 doses of mGlu_2_ and mGlu_3_ NAM (0, 3, 6, 12 mg/kg) were counterbalanced across test sessions and tested in both males and females using separate groups of rats for each drug tested (i.e., 4 separate groups). Final sample sizes for drug testing was as follows: male mGlu_2_-NAM (N=7) – 2 rats died from gavage during training/testing; female mGlu_2_-NAM (N=6) – 2 rats died from gavage during testing, male mGlu3-NAM (N=10), and female mGlu3-NAM (N=15).

## Data Analysis

For Experiment 1 (operant drug discrimination, Figure 1), response accuracy was evaluated as the percentage of alcohol-appropriate lever responding upon delivery of the first sucrose reinforcer (i.e., the first 10 lever responses). Full expression of the discriminative stimulus effects of alcohol was defined as ≥80% alcohol-appropriate responding. Partial substitution was defined as >40% and <80% alcohol-appropriate responding. No substitution was defined as <40% alcohol-appropriate responding. The response rate was calculated as lever responses per min and serves as a measure of motor output. For Experiment 2 (Pavlovian drug discrimination, Figure 2), the interoceptive effects of alcohol were evaluated as a discrimination score in response to the *first* light presentation (only 1 light presentation for test sessions). The discrimination score was calculated as [# of head entries into the liquid receptacle *during* the 15-s light presentation - # of head entries into the liquid receptacle in the 15-s *before* the light presentation]. This score served as a measure of behavioral activation in response to the cue and under these training conditions, head entries during the light presentations increase on alcohol, but not water sessions. As such, the discrimination score is a readout of the interoceptive effects of alcohol [30]. Locomotion was evaluated as beam breaks per min (locomotor rate).

**Fig. 1.**
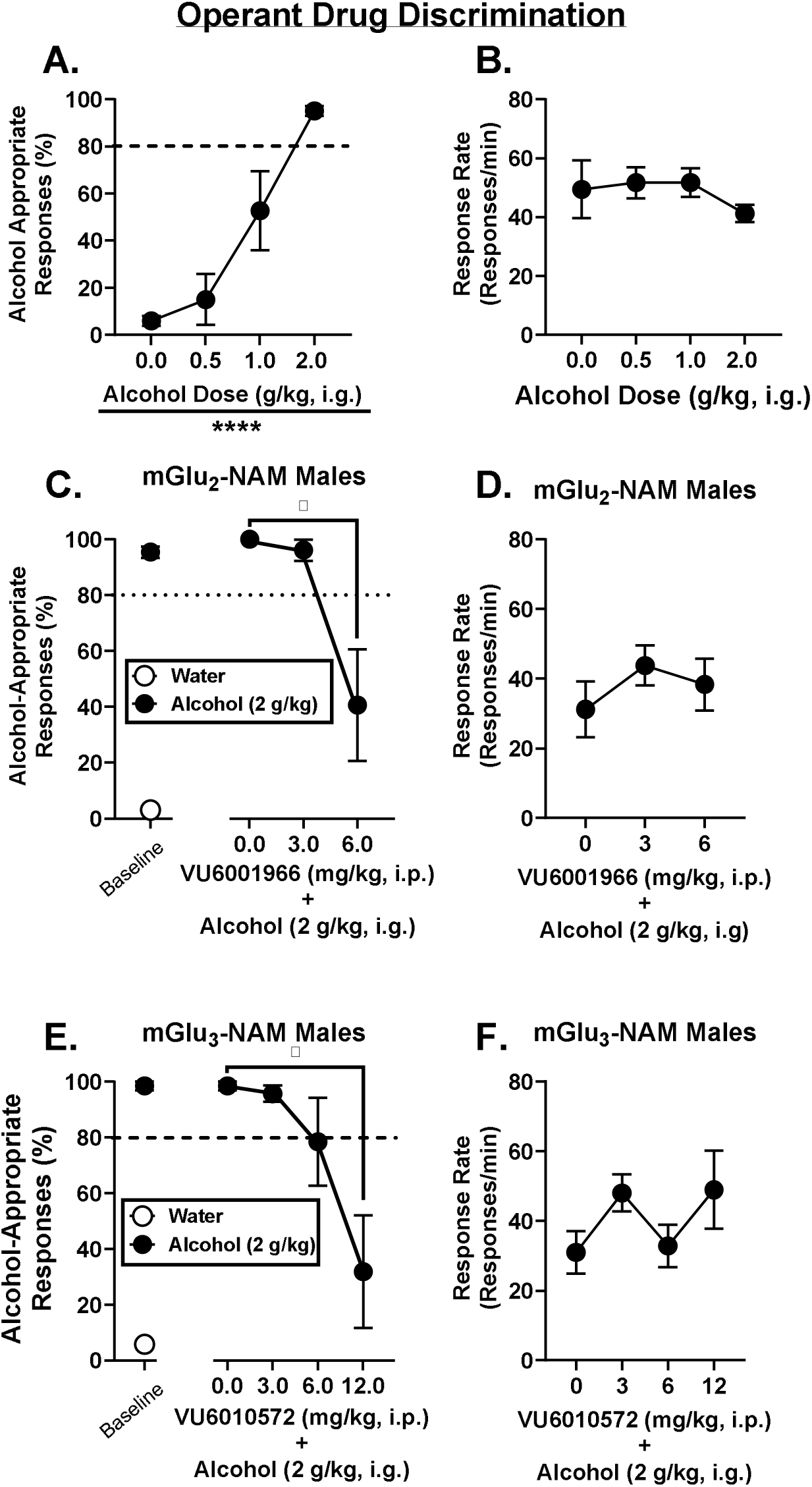
For operant drug discrimination, (A) alcohol dose response curve showed dose-dependent increases in alcohol-appropriate lever responding and (B) no effect of alcohol dose on response rate. (C) mGlu_2_-NAM VU6001966 attenuated alcohol-appropriate responding of 2.0 g/kg alcohol in males. (D) mGlu_2_-NAM did not affect response rate. (E) mGlu_3_-NAM VU6010572 attenuated alcohol-appropriate responding of 2.0 g/kg alcohol in males. (F) mGlu_3_-NAM did not affect response rate. Dotted lines at 80% indicate full substitution for the 2-g/kg alcohol-training dose. *p≤0.05. ****p<0.0001.

**Fig. 2.**
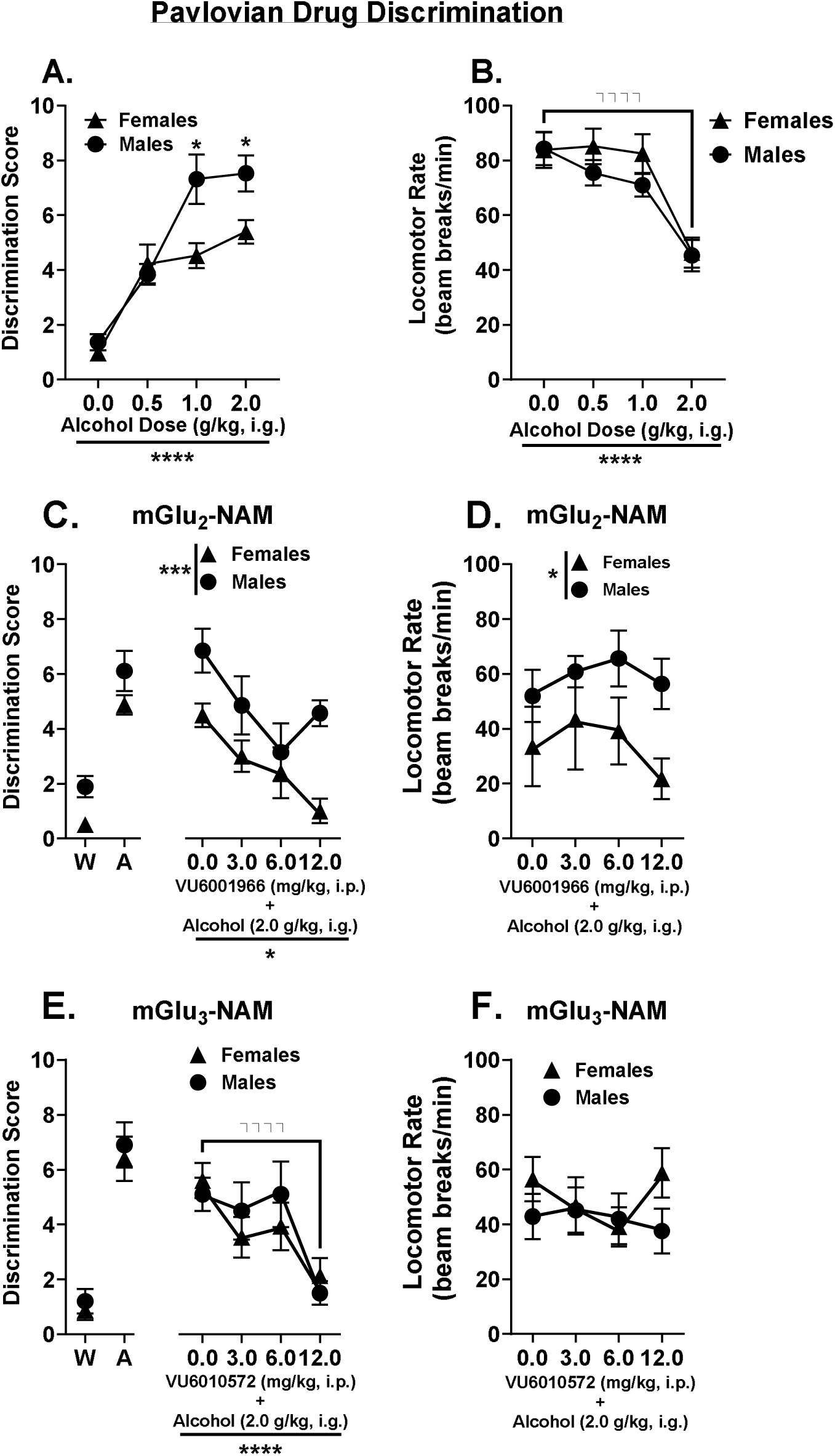
For Pavlovian drug discrimination, (A) the alcohol dose-response curve showed dose dependent increases in the discrimination score, with males showing higher discrimination scores than females. (B) Locomotor rate was attenuated at the 2.0 g/kg alcohol training dose. (C) mGlu_2_-NAM VU6001966 attenuated discrimination scores and there was a main effect of sex. (D) There was a main effect of sex for mGlu2-NAM locomotor rate, but no effect of drug dose. (E) The mGlu_3_-NAM VU6010572 attenuated discrimination scores and (F) did not affect locomotor rate. W-Water and A-Alcohol. *p≤0.05; ***p<0.001; ****p<0.0001.

The alcohol dose response curve for operant drug discrimination was analyzed using a repeated measures (RM) one-way ANOVA with Geisser-Greenhouse correction with alcohol dose as the within subjects factor. For mGlu NAM operant drug discrimination (Experiment 1; conducted in males only), a repeated measures (RM) one-way ANOVA with Geisser-Greenhouse correction was conducted with mGlu NAM drug dose as the within subjects factor (all rats received the 2.0 g/kg alcohol training dose during mGlu drug testing). Dunnett’s multiple comparisons test were conducted to compare each drug dose to vehicle. For Pavlovian discrimination (Experiment 2; in males and females), a RM-two-way ANOVA with Geisser-Greenhouse correction was used with alcohol dose as a within subjects factor and sex as a between subjects factor. For mGlu antagonism experiments, a RM-two-way ANOVA with Geisser-Greenhouse correction was used with mGlu NAM drug dose as a within-subjects factor and sex as a between subjects factor (mGlu NAM test days all rats received the alcohol training dose of 2.0 g/kg).

Effect sizes for all statistically significant results were calculated as eta squared (*η^2^*) for ANOVAs. For reference, a large effect size is defined as *η^2^* = 0.1379 or greater, a medium effect size is ^η*2*^ = 0.0588 or greater and a small effect size is *η^2^* = 0.009 or less [48]. P ≤ 0.05 was set for significance.

## Results

### Figure 1: mGlu_2_-NAM and mGlu_3_-NAM attenuate the interoceptive effects of alcohol using operant drug discrimination

For operant alcohol discrimination (conducted only in males), there was a main effect of alcohol dose for alcohol-appropriate lever responding (Fig. 1A; F (1.781, 14.25) = 20.13, p < 0.00001, *η^2^* = 0.60), showing dose-dependent increases with peak response and full substitution for the training dose of 2.0 g/kg alcohol. 1.0 g/kg alcohol produced partial substitution for the 2.0 g/kg training dose. There was no effect of alcohol dose on response rate (Fig. 1B). These data confirm stimulus control over lever selection behavior.

For reference, the final water and alcohol training session prior to testing is shown to the left of the x-axis break for the NAM experiment data. The mGlu_2_-NAM (VU6001966) significantly reduced alcohol-appropriate responding (F (1.02, 5.14) = 7.17, p = 0.04, *η^2^* = 0.52, Fig. 1C), with a reduction at 6 mg/kg compared to vehicle (p = 0.05). 6 mg/kg VU6001966 decreased alcohol-appropriate responding to levels considered “partial substitution” for alcohol - but note that the mean is at the cusp between “partial substitution” and “no substitution”. Alcohol-appropriate responding at 3 mg/kg mGlu_2_-NAM did not differ from vehicle and showed full substitution for the training dose. No effect of drug treatment was found on response rate (Fig. 1D). The mGlu_3_-NAM VU6010572 significantly reduced alcohol-appropriate responding (F (1.92, 9.61) = 6.72, p = 0.02, *η^2^* = 0.46, Fig. 1E), with a reduction at the 12 mg/kg dose relative to vehicle (p = 0.04), resulting in no substitution for the training dose. In contrast, alcohol-appropriate responding at 3 and 6 mg/kg did not differ from vehicle (p > 0.05), with 3 mg/kg showing full substitution and 6 mg/kg showing partial substitution. VU6010572 treatment did not affect response rate (Fig. 1F). These data indicate that the mGlu_2_-NAM and mGlu_3_-NAM both attenuate the interoceptive stimulus effects of 2.0 g/kg alcohol as measured by operant drug discrimination.

### Figure 2: mGlu_2_-NAM and mGlu_3_-NAM attenuate the interoceptive effects of alcohol in Pavlovian drug discrimination

For Pavlovian drug discrimination, discrimination scores increased as a function of alcohol dose (Fig. 2A, F (2.750, 110.0) = 47.57, p<0.0001, *η^2^* =0.39), confirming stimulus control. There was a main effect of sex (F (1, 40) = 6.661, p=0.0136, _η_*η^2^* =0.035) and an alcohol dose by sex interaction effect (F (3, 120) = 4.511, p=0.0049, *η^2^*=0.037). Males showed higher discrimination scores in response to 1.0 g/kg (p=0.01) and the 2.0 g/kg training dose (p=0.01). There was a main effect of alcohol dose on locomotor response in the Pavlovian discrimination paradigm (Fig. 2B, F (2.927, 117.1) = 28.48, p<0.0001, *η^2^*= 0.236), with 2.0 g/kg alcohol decreasing the locomotor rate compared to the other three doses (0.0, 0.5, 1.0 g/kg) tested in males and females combined (p<0.0001 for all main effects of dose).

For the NAM experiments, the final water and alcohol training session prior to testing is shown to the left of the x-axis break. For the mGlu2 NAM, there was also a main effect of drug dose (F (1.705, 18.76) = 6.041, p = 0.012; *η^2^* = 0.24), with a reduction in discrimination scores following 6 and 12 mg/kg compared to vehicle (p < 0.05). There was a main effect of sex (F (1, 11) = 21.14, p = 0.0008; *η^2^* = 0.19) with overall higher discrimination scores in the males (p<0.05). There was no sex by drug dose interaction. For mGlu_2_-NAM locomotion (Fig. 2D), there was no effect of drug dose, but there was a main effect of sex (F (1, 11) = 5.44, p = 0.04; *η^2^* = 0.17), with overall lower locomotor rate in females compared to males (p<0.05), and no drug dose by sex interaction.

Analysis of discrimination scores after pretreatment with the mGlu_3_-NAM (Fig. 2E), showed a main effect of drug dose (F (2.450, 56.36) = 9.319, p = 0.0001; *η^2^* = 0.18), with a significant reduction at the 12 mg/kg and 3 mg/kg doses compared to vehicle (p < 0.05 for 3 mg/kg; p < 0.0001 for 12 mg/kg). There was no main effect of sex, and no drug dose by sex interaction effect. The mGlu_3_-NAM did not affect locomotion (Fig. 2F) as there was no main effect of drug dose or sex, and no interaction effect. Together, these assessments replicate the findings from the operant discrimination experiment (Fig. 1) showing that the mGlu_2_-NAM and mGlu_3_-NAM attenuate the interoceptive stimulus effects of the 2.0 g/kg alcohol training dose.

## Discussion

These data show that systemic administration of both the mGlu_2_-NAM (VU6001966) and the mGlu_3_-NAM (VU6010572) attenuate the interoceptive effects of alcohol (2.0 g/kg training dose). These effects were observed using both operant and Pavlovian drug discrimination techniques and in both male and female rats. These are the first data to show the functional involvement of mGlu_2_ and mGlu_3_ separately in the expression of the interoceptive effects of alcohol in that negative allosteric modulation of mGlu_2_ or mGlu_3_ attenuated the interoceptive effects of alcohol.

Interestingly, in the Pavlovian discrimination experiment we also found that males showed greater sensitivity than females to the interoceptive effects of alcohol (2.0 g/kg training dose) when tested on the alcohol dose-response curve (0.0, 0.5, 1.0, 2.0 g/kg). One recent study from our lab using the same training procedures showed that males displayed slightly higher discrimination scores during a cumulative alcohol curve assessment when the training dose was 0.8 g/kg [43]. As such, male rats may be more sensitive to the interoceptive effects of alcohol compared to female rats, which may be more apparent at higher training doses of alcohol.

Previous work from our lab identified mGlu_2/3_ receptor involvement in the expression of the interoceptive effects of alcohol using an mGlu_2/3_ agonist in male rats [21]. In that study, systemic pretreatment with the mGlu_2/3_ agonist LY379268 blunted the interoceptive effects of 1.0 g/kg alcohol, and also produced increased neuronal activity in the amygdala subregions (central, basolateral, and lateral). Additionally, specifically targeting mGlu_2/3_ receptors in the central nucleus of the amygdala via microinjection of LY379268, but not the nucleus accumbens core blunted sensitivity to the interoceptive effects of 1 g/kg alcohol, confirming that the amygdala may be a key target region in mGlu_2/3_ receptor modulation of the interoceptive effects of alcohol. In another study, we found that systemic administration of the mGlu_2/3_ antagonist LY341495 also reduced sensitivity to the interoceptive effects of 1.0 g/kg alcohol [22]. That is, both the mGlu_2/3_ agonist and the antagonist produced the same behavioral effect in the operant alcohol discrimination task. This outcome raised the possibility that the mGlu_2/3_ agonist and antagonist system may produce unique interoceptive effects of their own that interfere with the expression of the alcohol interoceptive effects. The importance of the current work is that we were able to dissect the contribution of mGlu_2_ and mGlu_3_ separately using the NAMs. The finding that both mGlu_2_ and mGlu_3_ similarly produced a reduction in sensitivity to the interoceptive effects of 2.0 g/kg alcohol suggests a similar contribution of these receptors. It will be interesting for future work to investigate the brain regional contribution of each of these receptors.

There are some plausible explanations or mechanisms by which these NAMs attenuate the interoceptive effects of alcohol. Both the mGlu_2_ and mGlu_3_ NAMs have been shown to enhance c-Fos expression and increase excitatory currents on pyramidal prefrontal cortex neurons [41]. Alcohol exerts its interoceptive stimulus effects mainly through inhibition of the excitatory NMDA receptor and potentiation of the inhibitory GABA_A_ receptors [7, 32, 33, 39, 40, 49–51], which mediate much of the cellular firing behavior of neurons in the cortex. Therefore, the ability of mGlu_2_ and mGlu_3_ NAMs to block the interoceptive effects of alcohol may be due to the ability of these drugs to counteract the effects of alcohol on diminished cortical neuron excitation – i.e., through increasing neuronal excitation. Another interpretation is that mGlu_2_ and mGlu_3_ NAMs may produce their own interoceptive effects distinct from alcohol, which then interfere with the detection of the interoceptive stimulus of alcohol as previously mentioned [21, 22]. If mGlu_2_ and mGlu_3_ NAMs do produce their own distinct interoceptive stimulus effects, this may have implications for misuse and addiction liability of these drugs. Future experiments should be conducted to determine if these drugs can be trained as discriminative stimuli on their own, which would indicate potential for addiction liability. Note that these two explanations are not necessarily mutually exclusive and that there may be cortical circuitry that mediates interoceptive awareness and sensitivity in general rather than for any specific stimulus. Therefore, another explanation of these data is that mGlu_2_ and mGlu_3_ NAMs interfere with the detection, or encoding of interoceptive stimuli in general, and as such the interoceptive effects of alcohol are also blunted. Regardless of the precise mechanism(s), these studies demonstrate that systemically administered mGlu_2_ and mGlu_3_ NAMs may be used to attenuate the interoceptive stimulus effects of alcohol.

For the Pavlovian discrimination with the mGlu_2_-NAM (Fig. 2C), there appears to be a difference between males and females at the highest dose tested (12 mg/kg) such that males show higher discrimination scores compared to females. While a lack of a significant interaction effect precluded the post-hoc comparison, visual inspection of the data suggests a difference. This raises the possibility of a U-shaped drug effect in males – i.e., higher doses of mGlu_2_-NAM (> 6 mg/kg) may enhance, rather than attenuate, the interoceptive effects of alcohol in male rats. This discrepancy could be relevant to dosing of the mGlu_2_-NAM in males vs. females. Future experiments should test higher doses of mGlu_2_-NAM in both sexes to evaluate this hypothesis.

These are the first studies to demonstrate direct involvement of both mGlu_2_ and mGlu_3_ receptors in the interoceptive effects of alcohol. These results add to a growing literature on the involvement of mGlu_2_ and mGlu_3_ receptors in alcohol-related behaviors [9] and affective states [41]. These drugs have previously been shown to have acute anti-depressant-like behavioral effects in mice [26, 41] and produce prophylactic anxiolytic-like effects in rats [28]. As such, these drugs are promising for their potential in the treatment of affective disorders like depression and anxiety disorders. Because these disorders are highly co-morbid with AUD, and negative affect during withdrawal is an important part of AUD, these drugs may both alleviate the negative affect associated with AUD as well as attenuate the acute subjective effects of alcohol. Together, this combination of effects holds promise for improved pharmacotherapy of AUD.

## Contributions

**Ryan E. Tyler**: Conceptualization, Methodology, Formal Analysis, Investigation, Writing – Original Draft, Writing – Review and Editing, Visualization, Supervision

**Kalynn Van Voorhies**: Formal Analysis, Investigation

**Bruce E. Blough**: Resources

**Antonio Landavazo**: Resources

**Joyce Besheer**: Conceptualization, Methodology, Writing – Review and Editing, Resources, Supervision, Project Administration, Funding Acquisition

## Notes

Conflicts of interest: none

Funding: This work was supported in part by the National Institutes of Health [AA026537 and AA011605 (JB)] and by the Bowles Center for Alcohol Studies. RET was supported in part by NS007431 and AA029946.

### Competing Interest Statement

The authors have declared no competing interest.

### Summary of Updates

This version of the manuscript has been revised to include Pavlovian drug discrimination and testing in both male and female rats in addition to the original data using operant drug discrimination in male rats alone. Results were identical using either methodology and in both male and female rats.

